# Prenatal alcohol exposure disrupts hippocampal sharp-wave ripple-associated spike dynamics

**DOI:** 10.1101/2021.06.29.450435

**Authors:** Ryan E. Harvey, Laura E. Berkowitz, Daniel D. Savage, Benjamin J. Clark

## Abstract

Prenatal alcohol exposure (PAE) is among the most common developmental insults to the nervous system and is characterized by memory disruption. There is a pressing need to identify physiological alterations that help explain this memory impairment. Hippocampal sharp-wave ripples (SPW-Rs) are a compelling candidate for this purpose as they are the electrophysiological signatures of memory consolidation. We report that rats exposed to moderate prenatal alcohol display abnormalities restricted to SPW-R episodes that manifest as decreased recruitment of CA1 pyramidal cells and interneurons to SPW-R events, altered excitation during SPW-Rs, and decreased cell assembly activation rate. These differences observed at the single neuron and the population level may limit the ability of memory trace reactivation during SPW-Rs through the disruption of the intrinsic structure of cell sequences. Together, our results suggest that alterations in hippocampal SPW-R spike dynamics may underlie alcohol exposure-related memory deficits.

## Introduction

The dual-stage hypothesis for learning suggests that new associations are first stored within a network of hippocampal (HPC) synapses and are then propagated throughout the brain during high-frequency oscillatory bursts known as sharpwave ripples (SPW-Rs) (1–3). The “sharp-wave,” is a large amplitude excitatory wave prominent in the local field potentials of HPC subregion stratum radiatum, and the “ripple” is a high-frequency oscillation (80–250Hz) that rides atop the sharp-wave driven by synchronous firing among a large population of excitatory and inhibitory HPC neuronal ensembles. These events occur during moments of rest or sleep and are largely made up of pyramidal cells, including place cells, which fire in repeated sequences (4). This ensemble activity is generally known as reactivation, as firing sequences from active states are recapitulated during SPW-Rs and sometimes replay spatially coherent paths. Importantly, precise pyramidal cell-interneuron interactions have been found to support SPW-R generation as the silencing of pyramidal cells or activation of parvalbumin or somatostatin interneurons suppressed SPW-Rs (5). The HPC transmits this unique signal to the neocortex via the subiculum and retrospenial cortex (6–8). Together, this coordination is considered the underlying mechanism of memory consolidation via the strengthening of neuronal ensembles (9), as attempts to disrupt SPW-Rs have been shown to disrupt learning and memory (10–13). Likewise, Fernández-Ruiz et al. (14) found that spatial memory on an alternation task can be enhanced by optogenetically prolonging the duration of SPW-Rs. They concluded that this enhancement was likely due to the increased recruitment of new and diverse neurons to replay sequences during each SPW-R event. Correspondingly, the SPW-R associated spike dynamics during SPW-Rs are critical for the processes underlying learning and memory.

There is a large body of evidence that suggests that moderate amounts of alcohol exposure during fetal development (blood alcohol concentration 7-120 mg/dL) will negatively affect learning & memory via the disruption of synaptic and cellular networks (15–24). While the features of HPC CA3 sharp-waves are altered in-vitro slice preparations following a first human equivalent trimester ethanol exposure (25), how moderate PAE effects in-vivo SPW-Rs has yet to be explored. There is much evidence to suggest that PAE might alter SPW-Rs and replay. First, because moderate PAE during the first and second human equivalent trimester impairs spatial memory acquisition and retention (19,21–23,26,27), the underlying mechanism of memory consolidation may be disrupted. Secondly, moderate PAE reduces the number of HPC fast-spiking parvalbumin^+^ GABAergic interneurons (28) and disrupts HPC NMDA receptor-dependent long-term potentiation (29) indicating excitatory and inhibitory signaling is impaired. Broadly, PAE has been found to negatively affect GABAergic interneuron migration in multiple brain regions (discussed in (30)) which disrupts the balance between inhibitory/excitatory synaptic inputs later in life (31,32). Because SPW-Rs are supported by excitatory-inhibitory interactions (5), it might be expected that SPW-Rs would be altered following PAE. Lastly, functional NMDA-receptors are necessary for the formation of hippocampal sequences, largely composed of place cells, that replay spatial trajectories (33). The disruption of NMDA receptors (29) and place cell stability (17) observed following PAE may negatively impact the replay of spatial trajectories.

To explore how moderate PAE affects HPC SPW-Rs and replay, we recorded local field potential (LFP) and single units from hippocampal subregions CA1 and CA3 from 9 control and 8 PAE adult rats over several epochs as they rested on a holding pedestal, made laps on a linear track, and explored an open field (see Methods for details). SPW-R and replay events were detected while rats rested on the pedestal and while they paused to collect rewards during the two tasks, as SPW-Rs are most abundant during sleep or quiet wakefulness (34). Recordings were taken from hippocampal subregions CA1 & CA3, as SPW-Rs can be detected throughout the hippocampus (35,36).

## Results

### Features of sharp-wave ripples

To assess if PAE affected an inherent oscillatory aspect of SPW-R events, we measured the duration, frequency, amplitude, and SPW-R rate from CA1 & CA3 SPW-Rs. Data from all epoch types (pedestal, linear track, cylinder) were included as groups spent a similar amount of time in each epoch (*ps* ≥ 0.332) Fig. S1 and had similar numbers of unique epochs *χ*^2^(6,*N* = 1197) = 1.65, *p* = 0.95. As well, groups had a similar number of individual recording sessions (control: 172, PAE: 208 *χ*^2^(1, *N* = 380) = 3.41, *p* = 0.06). Because SPW-R duration is related to memory performance (14), we first measured the duration of each SPW-R event. Example SPW-Rs are shown in Fig. 1A-D. While all SPW-R features tested in CA1 were not significantly different between groups (*ps* ≥ 0.33, Fig. 1E-H), CA3 SPW-Rs from PAE rats had shorter durations (23ms median difference) compared with control CA3 SPW-Rs (*b* = −0.19, *SE* = 0.04, *p* = 0.00212, Fig. 1I). Likewise, the fraction of CA3 long duration SPW-Rs (≥ 100*ms*), which are indicative of diverse underling cell sequences (14), was decreased in the PAE group (Control 60%, PAE 44%, *b* =−0.57, *SE* = 0.17, *p* = 0.00089). However, other features of CA3 SPW-Rs such as SPW-R frequency, SPW-R amplitude, and SPW-R rate were not different between groups (ps ≥ 0.11, Fig. 1J-L). While the mechanisms that govern the duration of CA1 SPW-Rs seem unaffected by PAE, CA3 SPW-Rs are specifically affected.

**Figure 1.**
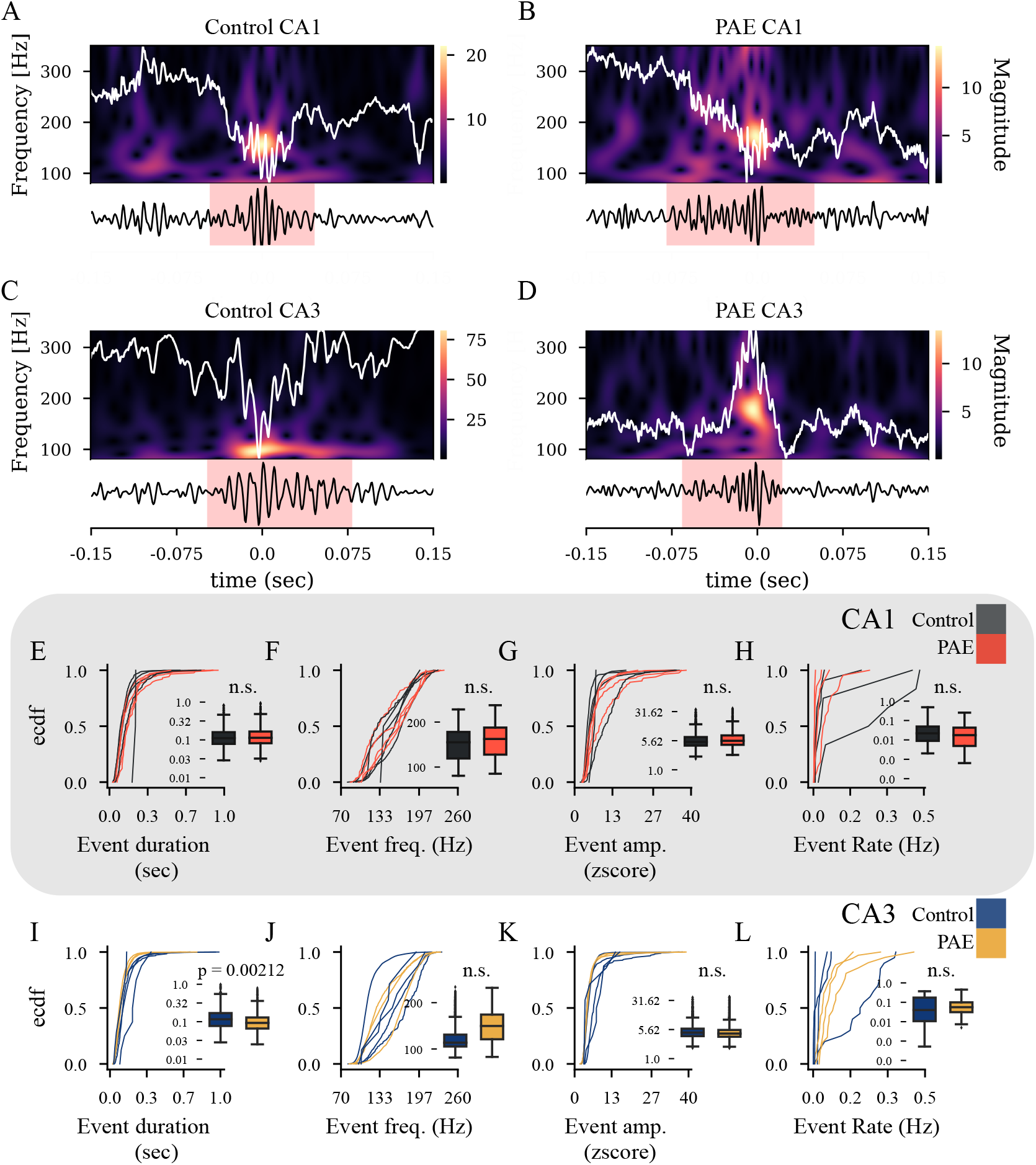
SPW-R examples and group comparisons. (A) Top: Shows a SPW-R example from control CA1. The white trace shows the wide-band (0-1250 Hz) signal which is superimposed on a time by frequency spectrogram. Bottom: Shows the band-pass filtered (80-250Hz) signal with a red shading that indicates the onset and offset of the SPW-R. (B-D) same as A but shows examples from PAE CA1, Control CA3, & PAE CA3. (E-L) Shows a comparison of SPW-R features between groups and HPC subregions. Each empirical cumulative density function (ecdf) represents data from a single rat. Inside each ecdf plot contains a standard box plot to highlight the group comparisons. (E-H) Data from CA1. (I-L) Data from CA3.

### CA1 cell recruitment to SPW-Rs is reduced following PAE

Hippocampal neurons fire synchronously during SPW-Rs (37,38), and longer-duration SPW-Rs were previously found to recruit a greater number of diverse hippocampal neurons (14). The diverse neurons that participate during long-duration SPW-Rs are thought to take part in ongoing neuronal sequences that assist in tasks that require higher memory demands. Because SPW-Rs from PAE rats were shorter, we sought to determine if neuronal recruitment during SPW-Rs was also affected. Because the disruption of interneurons has been found to reduce SPW-R duration (39), we investigated pyramidal cell and interneuron firing during SPW-R events.

We first classified pyramidal cells and interneurons using physiological characteristics such as waveform duration and autocorrelograms (Fig. S2 A-C). Proportions of CA1 & CA3 pyramidal cells were slightly lower in the PAE group (weighted mean proportions over sessions, CA1 control: 69%, CA1 PAE: 58%, CA3 control: 75%, CA3 PAE: 60%, Fig. S2 D-E). However, these proportions were not significantly different between groups (*ps* ≥ 0.061). To assess intrinsic firing properties of putative pyramidal cells and interneurons, we compared average firing rate, bursting propensity, and coefficient of variation II (CV2) (a measure of spike regularity (40)) between groups. No differences were identified for pyramidal and interneurons for either CA1 nor CA3 between groups in average firing rate (*ps* ≥ 0.30), bursting propensity (*ps* ≥ 0.114), or CV2 (*ps* ≥ 0.193). Together, these findings indicate that intrinsic properties on a single unit level are preserved following PAE.

Next, we investigated the relationship between neuronal spiking and SPW-Rs (Fig. 2) as individual units display strong recruitment into SPW-R events (41). Because CA3 SPW-Rs from PAE rats were shorter, it follows that there would be less diversity in the neurons participating in each SPW-R. To first investigate this, we assessed the relationship between the neuron’s within SPW-R rank order and baseline firing rate. Similar to previous work (14), the average rank order of a neuron’s within-SPW-R sequence was negatively associated with that neuron’s baseline firing rate (Fig. 2B) indicated that lower firing neurons are more likely to be recruited in the later portions of the sequence. CA1 pyramidal cells and interneurons and CA3 pyramidal cells from the PAE group had a weaker relationship between SPW-R rank order and baseline firing rate (CA1 pyramidal cells: *b* = 4.23, *SE* = 0.65, *p* = 1.86e – 10, CA1 interneurons: *b* = 5.55, *SE* = 1.01, *p* = 6.93e –08, CA3 pyramidal cells *b* = 1.26, *SE* = 0.60, *p* = 0.0368). This finding may reflect a disruption in the intrinsic structure of SPW-R sequences.

**Figure 2.**
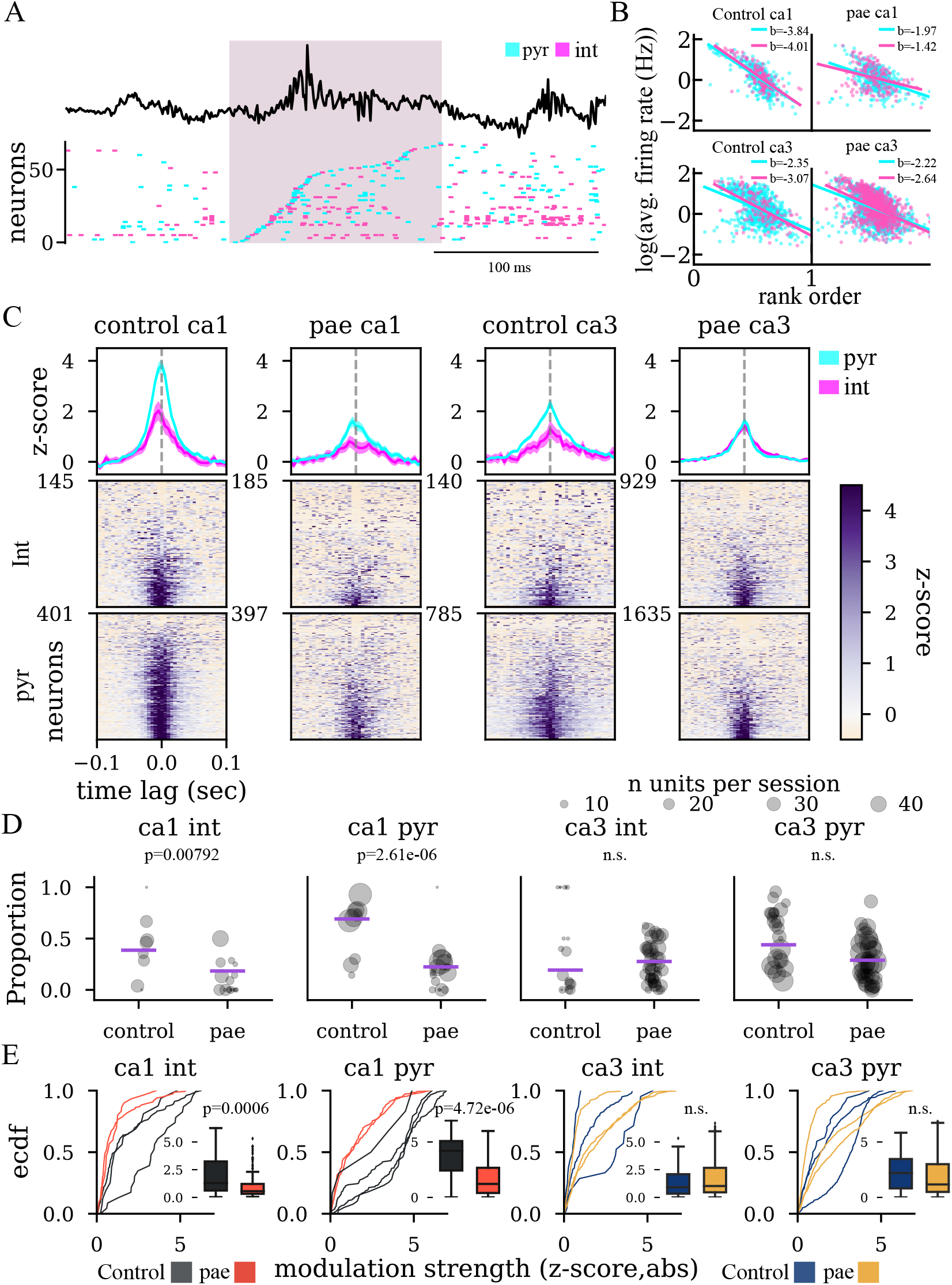
Temporal dynamics of individual units during SPW-Rs. (A) Example LFP trace centered around a SPW-R and neuron sequence below color-coded by cell type. Shaded segment of time represents SPW-R onset and offset. (B) Relationship between the average firing rate and the mean rank order in SPW-Rs. (C) Peri-SPW-R (±100*ms*) Z-scored firing rate plots for all putative pyramidal cells & interneurons. Top: average (with 95%confidence intervals) firing rate curves per group, region, and cell type shows a strong modulation of control cells to SPW-Rs. Bottom: All neurons organized by the time lag that maximized the firing rate of each unit. (D) Proportion of SPW-R modulated units. Group differences in proportions of classified units were compared with generalized linear models using rat and recording session as nesting factors. As such, data here is represented as a single point per recording session with the size of the point representative of the number of units recorded in that session. Weighted group averages are shown as purple lines over the scatted points. (E) Comparison of modulation strength between groups and HPC subregions. Each empirical cumulative density function (ecdf) represents data from a single rat. Inside each ecdf plot contains a standard box plot.

Next, we made peri-SPW-R rate plots for each cell and found that for a large proportion of cells, neuronal firing in CA1 & CA3 was time-locked to SPW-Rs with the greatest modulation observed in control CA1 pyramidal cells (Fig. 2C). Interestingly, fewer PAE CA1 pyramidal cells (PAE 22% vs. control 69% weighted average, *b* = −2.16, *SE* = 0.46, *p* = 2.61e –06) & interneurons (PAE 18% vs. control 37% weighted average, *b* = − 1.81, *SE* = 0.68, *p* = 0.00792) were significantly modulated with SPW-Rs (Fig. 2D), while the proportions of SPW-R modulated CA3 units was not significantly different between groups (*ps* ≥ 0.23). As well, we examined the modulation strength of each unit, defined as the absolute mean value of the z-scored center 3 bins of each peri-SPW-R tuning curve, and found decreased modulation strength of PAE CA1 interneurons (*b* =–1.15, *SE* = 0.29, *p* = 0.0006) and pyramidal cells (*b* = −2.13, *SE* = 0.37, *p* = 4.72e –06). Consistent with the number of modulated units, no significant differences in modulation strength were found in CA3 (*ps* ≥ 0.54).

Providing further evidence that PAE CA1 pyramidal cells and interneurons displayed decreased recruitment to SPW-Rs, we assessed participation at the single unit and population level (Fig. 3). The participation probability of individual units was significantly reduced in PAE CA1 interneurons (*b* = −0.69, *SE* = 0.33, *p* = 0.037) & pyramidal cells (*b* = −0.51, *SE* = 0.24, *p* = 0.034) while no significant differences were found in CA3 units (*ps* ≥ 0.15) (Fig. 3C). As the participation probability is positively related to the average firing rate (Fig. S3A), we examined average firing rate to rule out the possibility that the observed decrease in participation was simply due to lower firing rates of PAE units. As discussed above, average firing rates were not significantly different between groups (*ps* ≥ 0.30) (Fig. S3B). At the population level, we examined the fraction of units that were recruited for each SPW-R and found a smaller population of CA1 interneurons (*b* = −0.82, *SE* = 0.37, *p* = 0.0268) and pyramidal cells (*b* = −0.68, *SE* = 0.18, *p* = 0.000198) which participated in each ripple in the PAE group no significant differences in CA3 populations (*ps* ≥ 0.141 (Fig. 3B). Additionally, we examined SPW-R phase-locking, as neurons prefer to spike in specific SPW-R phases (41). We found that a greater proportion of PAE CA1 interneurons were phase-locked to a particular ripple phase (*b* = 1.65, *SE* = 0.51, *p* = 0.00111), however proportions from CA1 pyramidal cells and interneurons and pyramidal cells from CA3 were not significantly different (*ps* ≥ 0.52, Fig. S4). Together, there is sufficient evidence to conclude that, following PAE, interneurons and pyramidal cells from HPC CA1 are not being recruited into SPW-R events as strongly as units from the control group. However, because CA1 interneurons show a greater proportion of phase-locking, there could be compensatory mechanisms involved.

**Figure 3.**
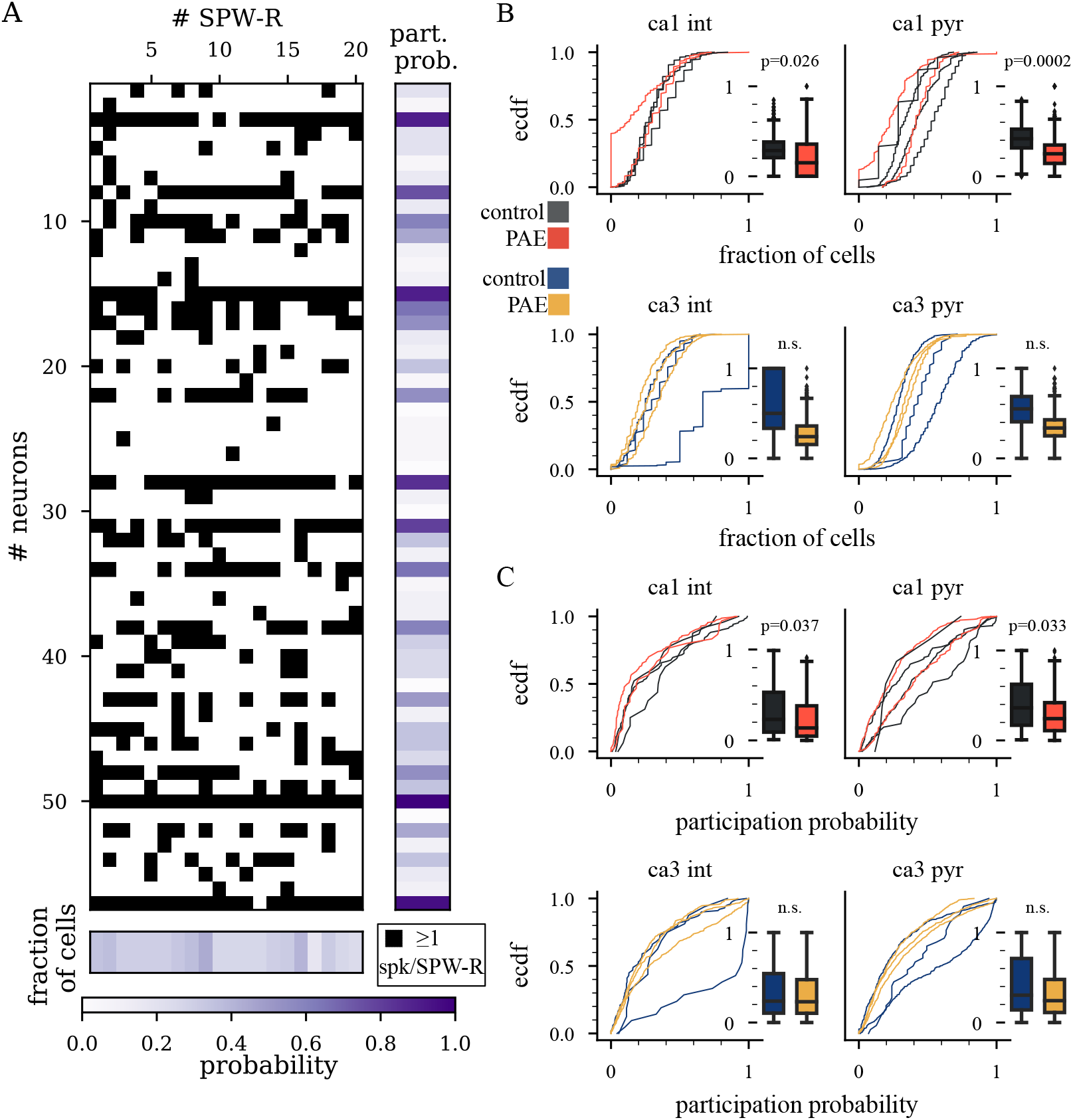
SPW-R participation. (A) Visualization of the participation analysis for a single recording session which produced the distributions compared in B and in C. The number of rows represents the number of neurons, and the black squares indicate whether the neuron fired at least one spike during a given SPW-R (columns). The right and bottom margins represent the participation probability for each neuron and the fraction of cells participating in a SPW-R respectively. Darker colors indicate a higher probability. (B) Distribution of the fraction of recorded neurons participating in each ripple calculated as the number of neurons that fired at least 1 spike during a SPW-R divided by the number of neurons recorded. (C) Participation probability. Group comparison of participation probability calculated as the number of SPW-Rs in which a unit fired at least 1 spike divided by the number of SPW-Rs that occurred in the session.

### Excitation and inhibition during SPW-Rs

Because of the decreased pyramidal cell and interneuron recruitment into SPW-R events, we next sought to characterize the roles of excitation and inhibition in local circuits during SPW-Rs through the quantification of pyramidal and interneuron population activity. Because SPW-Rs are supported by precise pyramidal cell-interneuron or excitation-inhibition dynamics (5,39,42–48), we hypothesized that the relative contribution of putative pyramidal cells and putative interneurons during SPW-Rs will be altered in moderate PAE.

To explore this hypothesis, we assessed the ongoing population activity of putative pyramidal cells and interneurons during SPW-Rs (Fig. 4A) as pyramidal cells are the primary excitatory, and interneurons are the primary inhibitory cell type in the mammalian brain. Similar to single units, pyramidal cell and interneuron population activity increased during SPW-Rs, which allowed the comparison of their relative contribution to each SPW-R event. We found that the control group’s CA1 SPW-Rs were more dominated by pyramidal cell activity whereas the relative contribution of pyramidal cell-interneurons to PAE CA1 SPW-Rs was more balanced (*b* = −71.30, *SE* = 0.51, *p* = 0.0119, Fig. 4B). This finding is supported by investigating the within SPW-Rs peak population strength of pyramidal cells (Fig. 4C) and interneurons (Fig. 4D). Population activity of pyramidal cells during CA1 SPW-Rs was significantly higher in the control group (*b* = −0.11, *SE* = 0.03, *p* = 0.00363), while interneuron population activity was not significantly different between the groups (*p* = 0.875). Consistent with our previous finding of decreased CA1 pyramidal cell and interneuron recruitment into SPW-Rs, pyramidal cell–interneuron dynamics in CA3 were unaffected across these features (Fig. 4E-G, *ps* ≥ 0.0573). Together, these findings suggest that CA1 network activity is less biased towards excitation during SPW-Rs in rats with PAE. This decreased excitation may explain the previously discussed CA1 specific reductions in cell recruitment.

**Figure 4.**
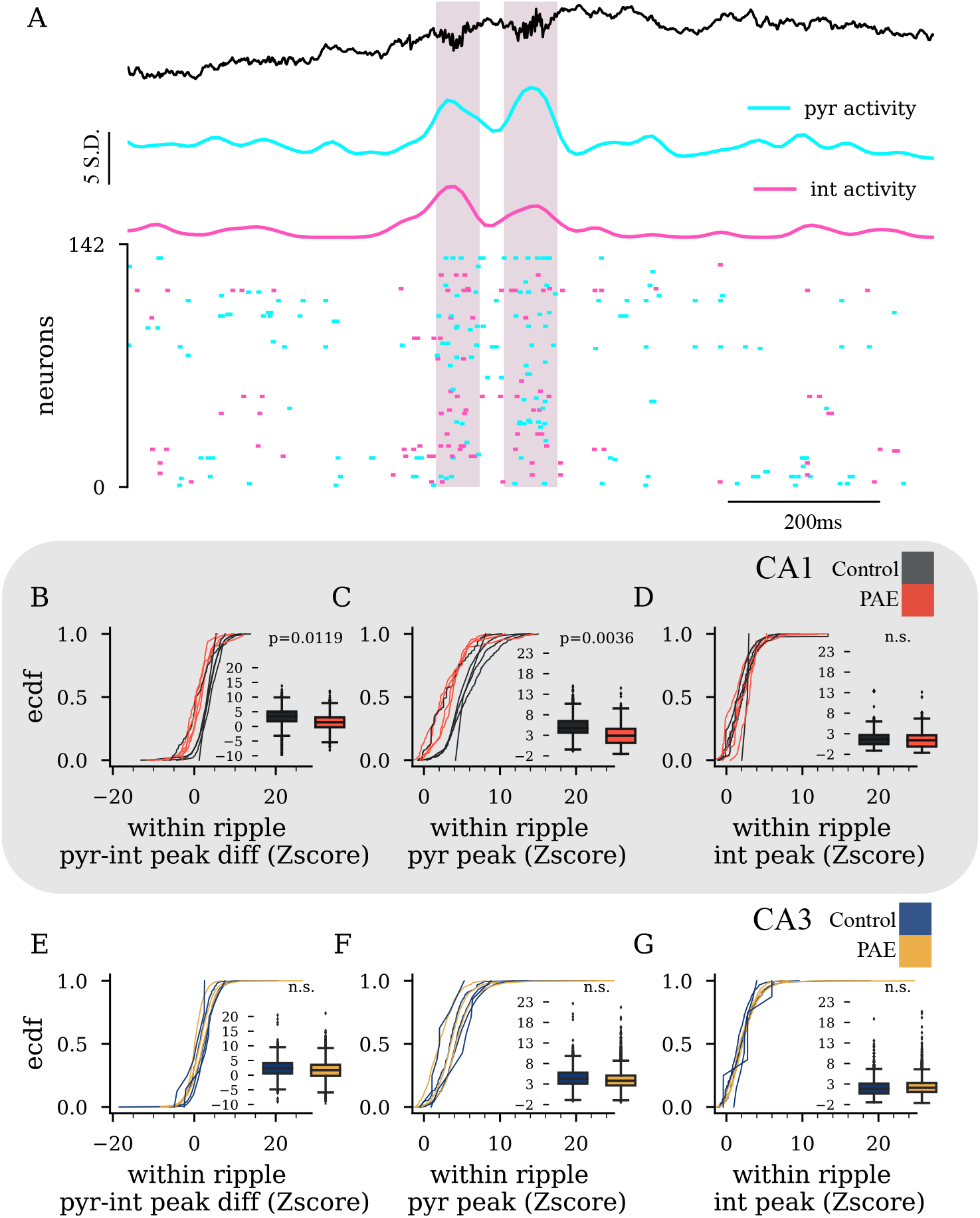
The relative contribution of excitatory and inhibitory signals during SPW-Rs. (A) Example demonstrating population activity over time and its association to SPW-Rs and spiking activity. Top: raw LFP with shading to denote SPW-R epochs. Middle: concurrent population activity from pyramidal cells (pyr) and interneurons (int). Bottom: Raster plot of individual units color coded by their cell classification. (B) Comparison of the within-ripple pyramidal-interneuron peak difference (pyramidal population activity peak subtracted by interneuron population activity peak for each SPW-R epoch). Note that the control group’s scores are larger indicating that, in the control group, pyramidal cell activity was larger relative to inhibitory activity during SPW-Rs, whereas pyramidal and interneuron activity was more similar in the PAE group. (C) Comparison of the within ripple pyramidal peak activity (peak value of population activity within each SPW-R epoch). Note that scores from the PAE group as significantly decreased. (D) Comparison of the within ripple interneuron peak activity (peak value of population activity within each SPW-R epoch). (E-G) Same comparisons as B-D, but for CA3. Note that comparisons within CA3 were not significantly different.

### Hippocampal spatial replay is unaffected by PAE

Because the CA1 unit activity during SPW-Rs is decreased in rats with PAE, it was thought that the replay of spatial content during SPW-R events would likely be altered. To investigate hippocampal replay, we used a Bayesian approach (49) where we constructed average firing rate-maps for each recorded neuron to estimate the probability distribution of the animal’s position on the linear track over time during SPW-R events (50,51). In order to be confident in our estimation of spatial information during replay events, we first confirmed that the quality of decoding was not significantly different between groups (*ps* ≥ 0.125, median decoding error: CA1 control 6.12 cm, CA1 PAE 5.51 cm; CA3 control 10.90 cm, CA3 PAE 7.88 cm). To maximize the number of detected replay events, we investigated epochs of increased multi-unit synchronous activity (MUA) (50).

Diverging from the CA1 differences in SPW-R recruitment, awake replay of spatial trajectories in CA1 & CA3 was un-affected by PAE. Specifically, the decoded spatial trajectories (Fig. 5A) during MUA events did not significantly differ in their Bayesian trajectory scores (sum of the probabilities under the line of fit), overall trajectory distance, trajectory speed, or bin-to-bin spatial step size (*ps* ≥ 0.089) (Fig. 5B-I). As well, the proportion of events demonstrating forward and reverse trajectories, relative to the rat’s current position, was not significantly different between groups (*ps* ≥ 0.927). Finally, the proportions of significant replay events out of all valid candidate events were found in similar proportions to previous studies (50,52–56) and were not significantly different between groups in either CA1 nor CA3 (*ps* ≥ 0.099, session mean proportion: CA1 control 15%, CA1 PAE 19%, CA3 control 32%, CA3 PAE 17%). Together, the evidence suggests that PAE does not affect awake hippocampal replay of spatial trajectories.

**Figure 5.**
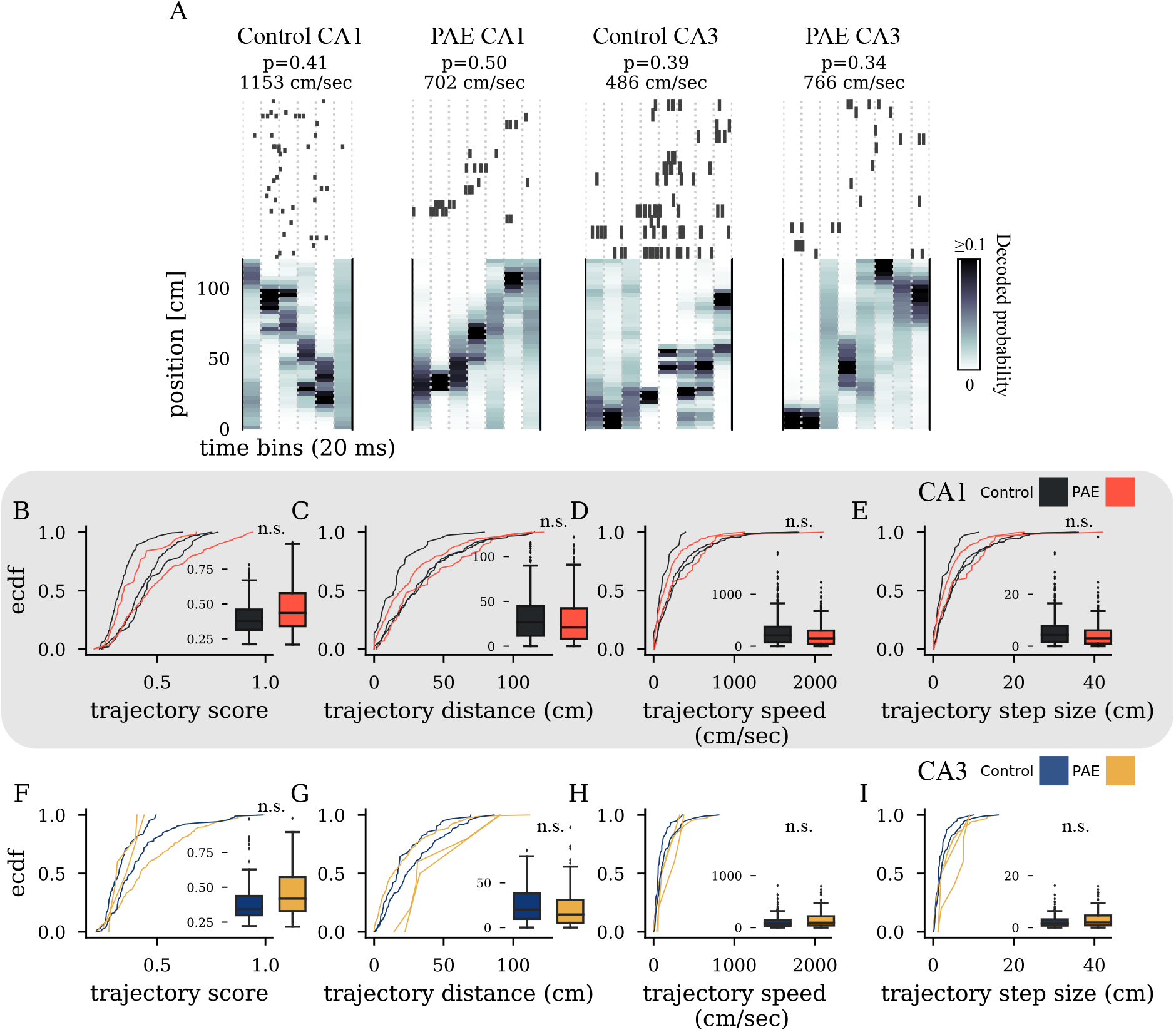
Similar replay events between groups. (A) Raster plot of spikes during example MUA events sorted by peak tuning curve position (top) and their associated time-by-position probability derived from Bayesian decoding of position (bottom). Color shows decoded probability from 0 to ≥ 0.1. The summed probability under the fit line (p) and speed (cm/sec) of event are indicated above the raster plots. (B) Replay trajectory score for control (gray) and PAE (red). (C) Distance of the linear track spanned by each replay event. (D) Speed in cm/sec of replayed trajectory from starting position to end position for all replay events. (E) Mean spatial step or jump in centimeters for decoded positions in adjacent temporal bins. (F-I) Same as B-E, but for HPC CA3.

### CA1 cell assembly activation rate is diminished by PAE

While the awake replay of spatial trajectories was unaffected by PAE, we sought to investigate sequence activity more broadly as changes observed in single-unit activity during SPW-R events may affect the precise organization of activity across cell networks. To investigate network activity, we used a combination of principal component and independent component analyses to identify and monitor coactivated groups of neurons often referred to as cell assemblies or ensembles (57,58) (Fig. 6A). Assemblies were identified using data from an entire recording session and then investigated during SPW-R epochs. CA1 & CA3 assembly strengths, measured by their peak activation values during SPW-R epochs, were not significantly different between the groups (*ps* ≥ 0.15, Fig. 6B,E). As well, the number of units with significant contribution to each assembly was not affected by PAE (*ps* ≥ 0.156). However, the rate at which CA1 assemblies were activated during SPW-R epochs was significantly higher in the control group (*b* = −0.44,*SE* = 0.14,*p* = 0.0035, Fig. 6C). Similarly, the proportion of CA1 SPW-Rs that displayed at least one assembly activation was also higher in the control group (*b* = −1.07, *SE* = 0.26,*p* = 5e –05, Fig. 6D). Together, the findings suggest that PAE partially disrupts the ability of CA1 cells to precisely coordinate their firing in a way to support network activity during SPW-R events, which may in turn interrupt the consolidation of memories.

**Figure 6.**
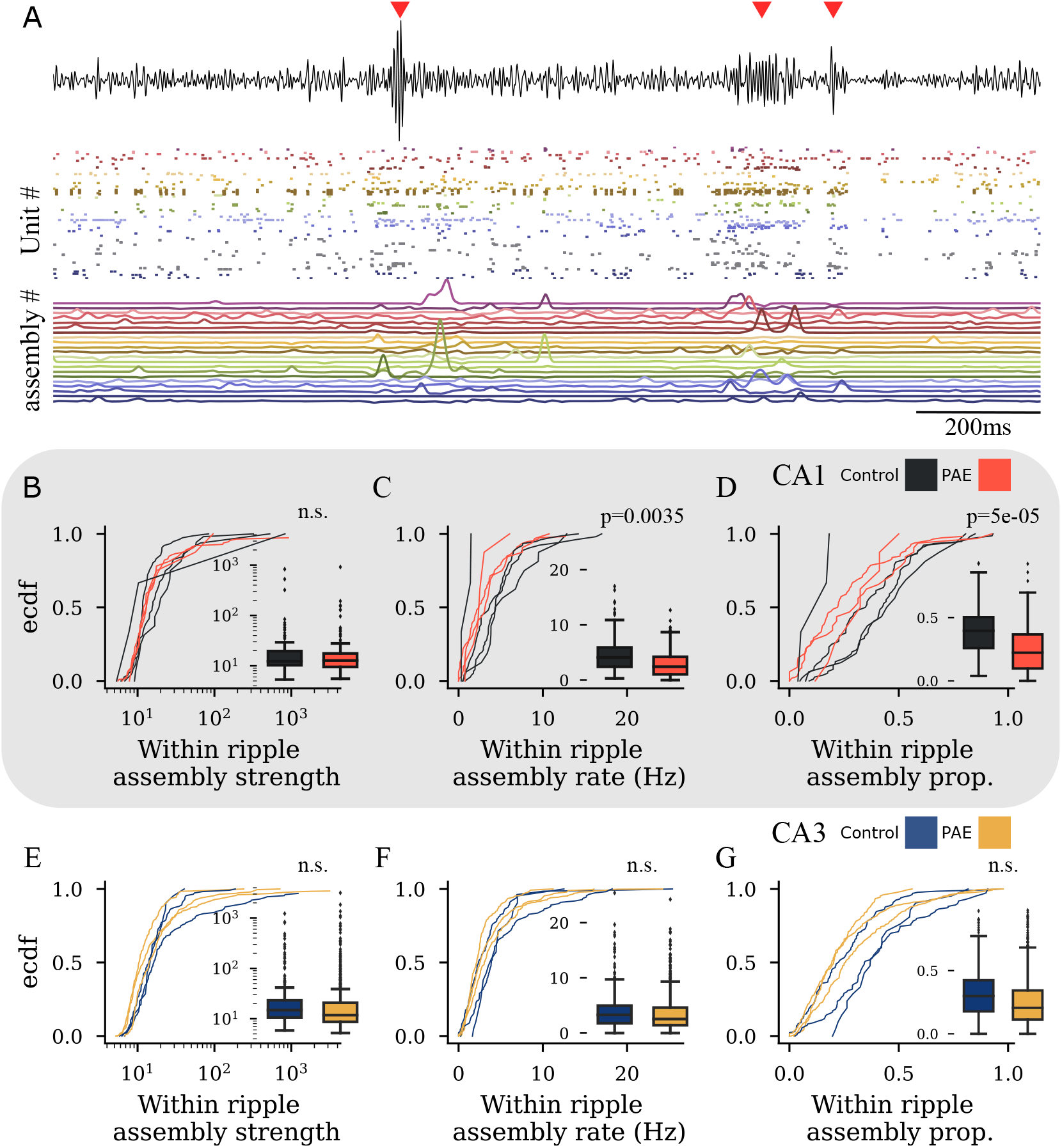
Cell assembly properties. (A) Example demonstrating assembly activity over time and its association to SPW-Rs and spiking activity. Top: filtered LFP with red triangles to denote SPW-R epochs. Middle: concurrent spiking activity color-coded by their associated assembly. Units not significantly associated with any assembly are shown in light gray. Bottom: Assembly expression strength for each assembly over time. (B-D) Group comparison of within SPW-R assembly strength, rate, and proportion of SPW-Rs that contained significant assembly peaks (*R* > 5) for CA1. (E-G) Same as B-D but for CA3. Each empirical cumulative density function (ecdf) represents data from a single rat. Inside each ecdf plot contains a standard box plot.

## Discussion

The central theory behind this study was that SPW-Rs are vital for memory consolidation through the strengthening of neuronal ensembles (1–3,9). Because learning and memory are impaired following prenatal alcohol exposure, we sought to document changes to SPW-R characteristics in a model of prenatal exposure.

There is a large body of evidence that has identified SPW-Rs as a critical mechanism in learning and memory (11,13,50,53,54,58–66). Neurological disorders that are characterized by deficits in learning and memory, such as PAE, may specifically disrupt the underlying mechanisms that produce SPW-R events. Our characterization of the SPW-R events is consistent with this.

Here we report that rats exposed to moderate prenatal alcohol display abnormalities restricted to SPW-R episodes that manifest as decreased CA3 SPW-R durations, decreased recruitment of CA1 pyramidal cells and interneurons to SPW-Rs, and decreased cell assembly activation rate. Awake hippocampal replay sequences of spatial trajectories were unaffected by PAE. Our results suggest that alterations in hippocampal SPW-R associated spike dynamics may underlie alcohol exposure-related memory deficits.

Many characteristic oscillatory features of HPC SPW-Rs were similar between groups, such as frequency, amplitude, and rate. It is only in CA3 we found that SPW-R durations were slightly shorter following PAE (Fig 1I). Moderate PAE reduces the number of HPC fast-spiking parvalbumin^+^GABAergic interneurons (28) which are critical for sustaining long-duration SPW-Rs (39). Thus, the lack of CA3 GABAergic interneurons in this model may underlie our finding of reduced CA3 SPW-R duration. This finding contrasts with a previous study by Krawszyk et al. (25) that found longer sharp-wave durations of CA3 ripples following PAE. Many reasons exist that may account for this discrepancy, such as the difference in the timing and dosage of ethanol, the difference in in-vivo vs. in-vitro SPW-Rs, or differences between SPW-Rs in rats and mice (16,67–69).

Although only subtle differences were observed in the oscillatory component of SPW-Rs, changes in the underlying neuronal activity were evident. Specifically, the recruitment of CA1 pyramidal cells and interneurons to SPW-Rs was reduced (Fig 2 & 3). This reduction was observed in the number of cells that significantly changed their firing (Fig 2D) as well as the magnitude of firing change during SPW-Rs (Fig 2E). Because the recruitment of HPC neurons to SPW-Rs is critical for learning and correct recall in spatial memory tasks (14), the decrease in recruitment limits the ability of memory trace reactivation during SPW-Rs. This deficit may underlie some reported issues with learning and memory such as the ability to rapidly acquire and retain spatial information over multiple days (16,26,27,70).

Because of the decrease in cell recruitment into SPW-R events, we sought to examine the role of excitation and inhibition of local circuits. Consistent with previous studies examining the HPC and cortical regions following moderate PAE (25,31,32), the current study found an excitatory-inhibitory imbalance in the HPC following PAE. Specifically, there was decreased CA1 pyramidal cell population activity relative to interneuron activity during SPW-Rs, indicating a reduction of local excitation. While we found no difference in CA3, Krawczyk et al. (25) found an excitatory-inhibitory imbalance in favor of CA3 excitation following PAE.

This contradiction may be due to their higher ethanol exposure which produced a peak BAC of ~ 120 mg/dl in contrast to ours which produced a peak BAC used ~ 60 mg/dl. Alternatively, the difference may be due to their patch-clamp paired-pulse approach. Being so, it is important to highlight the difference in the approach used here compared to previous work. Previous studies have examined excitatory/inhibitory balance during SPW-Rs via the recording of excitatory and inhibitory postsynaptic currents (EPSCs and IPSCs) (44,71), as recording EPSCs and IPSCs is a more direct measure of excitatory/inhibitory balance. However, because pyramidal cells are the primary excitatory, and interneurons are the primary inhibitory cell type in the mammalian brain, their population activity analyzed here should constitute an accurate estimation of the relative excitation and inhibition of local circuits.

Despite alterations to CA1 cell recruitment and imbalance in local excitation/inhibition dynamics, hippocampal replay of spatial trajectories was found to not differ following PAE (Fig 5). However, consistent with prior literature, the minority (~ 5 —20%) of SPW-Rs contain significant spatial trajectories (50,52–56). The replay tested here only concerned the spatial content of SPW-Rs, while the differences in recruitment and assembly activity might reflect non-spatial domains of replay. The issue of detecting the replay of non-spatial features of experience is discussed in Tingley & Peyrache (72). One would need to include every aspect of the animal’s experience in a model in order to fully examine what a candidate replay event is truly ‘replaying.’ For example, Oliva et al. (73) discovered a subset of SPW-Rs from HPC CA2 that reactivate and consolidate social memories. As such, while spatial replay is unaffected by PAE, the replay of other dimensions of the animal’s experience might be affected.

We next asked whether co-active assembles of neurons were similarly reactivated during SPW-Rs following PAE (Fig 6). While the strength of reactivation was similar between groups, the rate at which CA1 neurons were reactivated during SPW-Rs was decreased compared with control CA1 neurons. This decrease suggests that HPC replay might be incomplete, and aspects supported by computations at this level, such as learning and memory (11,13,50,53,54,58–66), might be perturbed. The observed effects here may underlie some alcohol exposure-related memory deficits such as impaired spatial working memory for locations observed in the Morris water maze (19,21,74) or post-acquisition retention impairments (27). It would be important to test whether alterations in SPW-R activity can predict memory performance in a within-study design similar to Jones et al. (75). Doing so would be a more direct examination of how the alterations shown here can influence behavioral performance.

Here we report robust changes in CA1 but not CA3 at the single cell level, demonstrated by a decrease in participation probability and lack of strong firing with SPW-Rs, and at the population level, demonstrated by the reduced activation rate during SPW-Rs. These findings are likely an outcome of changes on the network level as intrinsic cell properties are preserved following PAE. Instead, these changes may be due to deficits in local CA1 pyramidal cell-interneuron interactions as evident by a decrease in pyramidal cell population activity highlighted in Fig4. It is likely that this difference in excitation and inhibition throughout the CA1 network would disrupt underlying SPW-R spiking, as SPW-Rs generation and underlying spiking dynamics are supported by precise pyramidal cell–interneuron interactions (5,39,42–48).

There are several unanswered questions remaining. For instance, recordings were not taken while rats slept. Memory consolidation via SPW-Rs has been classically described to occur during non-REM sleep (discussed in Buzsáki, (4)). Children and infants with FASD have clinically significant sleep problems such as fragmented sleep, disordered breathing during sleep, increased sleep anxiety, frequent awakenings, and reduced total sleep time (76–79). These sleep problems undoubtedly disturb non-REM sleep, which in-turn may alter the occurrence of SPW-Rs during sleep. It would be a benefit for future studies to investigate the effects of PAE on SPW-Rs and related neuronal activity during post-task sleep.

Another open question is how HPC subregions interact during SPW-Rs. In this current work, while subregions were separated, simultaneous layer-specific recordings were not collected. The use of two-dimensional silicon probe arrays, such as used by Oliva et al. (35), would allow the simultaneous investigation of the propagation of SPW-Rs across HPC CA1, CA2, CA3, and DG subregions. This added information would better inform how the activity spreads throughout the subregions throughout the generation of a SPW-R, which would allow us to address hypotheses about the interaction between regions. Because there is evidence of altered long-term potentiation in the entorhinal cortex → dentate gyrus circuit following PAE (29,80–88), the dentate gyrus → CA3 circuit following postnatal exposure (89), and the CA3 → CA1 circuit (90–93), it is hypothesized that the precise sequence of SPW-R propagation throughout the subregions (discussed in detail in Oliva et al. (35)) may be altered following PAE. For this reason, the decrease in CA3 SPW-R duration may be due to the lack of CA1 recruitment as co-recorded CA3 ripples can occur following CA1 SPW-Rs (35,94). Likewise, the impact of HPC SPW-Rs on downstream regions of CA1 such as the subiculum and retrosplenial cortex (6) is likely to be reduced, which indicates that the ability of the hippocampus to convey neuronal content to the neocortex during SPW-Rs may be severely limited following PAE.

### Conclusions

This study adds to the extensive literature detailing the effects of prenatal alcohol exposure, which is among the most common developmental insults to the nervous system. It is our hope that this study helps to connect the many studies separately examining the effects of PAE on memory performance and synaptic and cellular properties. To do this, we examined the functional mechanisms underlying memory consolidation via the phenomena of hippocampal sharp-wave ripples and found robust alterations in the ripple-associated spike dynamics. These differences may underlie alcohol exposure-related memory deficits. We are optimistic that this paradigm may provide a novel testing platform for future therapeutic interventions.

## ACKNOWLEDGEMENTS

The authors thank Dr. Suzy Davies, and Dr. Jennifer Wagner for supervising the moderate PAE paradigm and Andre Moezzi, Nicole Graham, Kiana Lujan, Ella Rappaport, and Chloe Puglisi from the UNM Department of Neurosciences as well as Gilbert Borunda and Sean Bilberry from the UNM Department of Psychology for their assistance with the rodent husbandry procedures. We would also like to thank Azahara Oliva and Antonio Fernández-Ruiz for helpful comments on the manuscript. The Clark and Savage Labs were supported by National Institute on Alcohol Abuse and Alcoholism grants R21 AA024983, P50 AA022534, T32 AA014127-15 and a Grice Research Enhancement Award from The UNM Department of Psychology.

## AUTHOR CONTRIBUTIONS

R.E.H., D.D.S., and B.J.C. conceptualized experiments. R.E.H. designed the study and conceptualized the data analysis. B.J.C. contributed lab resources and supervision. D.D.S. created the animal model and directed the production of experimental offspring. R.E.H. collected, analyzed, and conducted statistical analyses on the data with help from L.E.B. R.E.H. wrote the initial draft of the paper with B.J.C., D.D.S., and L.E.B. providing review and editing.

## Resource availability

### Lead contact

Further information, materials, and protocols used in the manuscript will be made available upon request to the lead contact authors, Ryan E. Harvey or Benjamin J. Clark.

### Data and Code Availability

The data-set used for this study is available at OSF: https://doi.org/10.17605/OSF.IO/PM89Y. Custom code is available at https://github.com/ryanharvey1/ephys_tools, https://github.com/ryanharvey1/ripple_analyses, and https://github.com/ryanharvey1/cell_assembly_replay.

## Materials and methods

### Experimental Model and Subject Details

#### Subjects

Subjects were 17 male Long-Evans rats (4—6 months old) obtained from the University of New Mexico Health Sciences Center Animal Resource Facility. Data from all rats contributed to a previous study (17) where breeding, ethanol consumption, behavioral training, surgical procedures, and recording procedures are described in detail. All procedures were approved by the Institutional Animal Care and Use Committee of either the main campus or Health Sciences Center at the University of New Mexico.

#### Breeding and voluntary ethanol consumption during gestation

Three to four-month-old female breeders (Harlan Industries, Indianapolis, IN) were exposed to a voluntary ethanol drinking paradigm described in detail in previous studies (17,21,22,26–28,74,87,95–99). In brief, female rats were provided 0.066% (w/v) saccharin in water for 4h each day. On days 1—2, the saccharin water contained 0% ethanol, on days 3—4 saccharin water contained 2.5% ethanol (v/v), and on day 5 and thereafter, saccharin water contained 5% ethanol. After two weeks of daily ethanol consumption, rats that drank at levels outside one standard deviation of the entire group mean were removed, and the remaining rats were randomly assigned to either a saccharin control or 5% ethanol drinking groups.

Female rats were matched with a male breeder rat until pregnancy was verified. Beginning on gestational day 1, the rat dams were given access to saccharin (Sigma Life Sciences, St. Louis, Missouri) water containing either 0% (v/v) or 5% (v/v) ethanol (Koptec, King of Prussia, Pennsylvania) for four hours a day. During gestation and including the 4h ethanol/saccharin drinking period, rats were provided with ad libitum water and rat chow (Teklad global soy protein-free extruded food 2920). The daily mean ethanol consumption throughout pregnancy was 1.96 ±0.14 g/kg) and did not vary significantly during each of the three weeks of gestation. Daily ethanol consumption ended at birth. Offspring were aged to 4—6 months of age before behavioral training and electrophysiological recordings took place.

#### Behavioral training

Each animal was exposed to each maze for several days. On the linear track, animals were trained to continuously alternate between the two ends. Stop locations were rewarded with 1/8 piece of a fruit-loop if animals completed a successful lap without stopping or alternating the incorrect direction. This training continued until 40 laps could be completed within 15 minutes. For the cylinder, animals were trained to randomly forage for fruit-loop pieces that were semi-randomly dropped into the environment until all locations were sufficiently sampled. Experimenters were aware of each animal’s group affiliation (PAE or control) upon initiation of behavioral training.

### Method Details

#### Recordings

Rats were implanted with either an 8 tetrode bundle or 16 independently moveable tetrodes targeting hippocampal subregions. Single units and local field potentials were recorded while rats collected food pellets on a linear track (120 cm by 9 cm), open cylinder environment (diameter: 76.5 cm), or resting on a small pedestal. Pedestal rest epochs lasted around 2 minutes on average.

The electrode assembly was connected to a multi-channel impedance matching unity gain preamplifier headstage. The output was routed to a data acquisition system with 64 digitally programmable differential amplifiers (Neuralynx, Tucson, AZ, USA). Spike waveforms above a threshold of 30–40 *μ*V were time-stamped, digitized at 32 kHz, and later imported into MClust for spike sorting. Continuous wide-band signals from each channel were collected for sessions which were spike sorted with Kilosort2. The rat’s position was tracked and sampled at 30 Hz by recording the position of light-emitting diodes that were placed above the head.

### Quantification and Statistical Analysis

#### Data analysis

Data analysis was performed by processing position, LFP, and spike data with custom Matlab code https://github.com/ryanharvey1/ephys_tools and further processing the data with custom-written Python code. Custom ripple analyses: https://github.com/ryanharvey1/ripple_analyses. Custom cell assembly and replay analyses: https://github.com/ryanharvey1/cell_assembly_replay. Many computations were carried out using NumPy (100). Statistical analysis was performed in R. Data visualization used Matplotlib (101), Seaborn (102), and Nelpy (103).

#### Spike sorting

Spike sorting was performed first by using unsupervised methods from klustakwik http://klustakwik.sourceforge.net/ or Kilosort2 https://github.com/MouseLand/Kilosort2 with manual refinement conducted in MClust https://github.com/adredish/MClust-Spike-Sorting-Toolbox or Phy https://github.com/cortex-lab/phy. The inspection of autocorrelation and cross-correlation functions were used as single unit identification criteria for both manual refinement methods.

#### Pyramidal and interneuron classification

Cells were classified into putative pyramidal and interneurons based on their waveform duration obtained using a wavelet transform. Waveforms were obtained by first filtering the wideband signal between 600 and 9000 Hz, and then taking a 1ms window relative to the spike timestamp. Waveform duration was estimated using the inverse of the maximum power associated frequency in its power spectrum; obtained from a wavelet transform (104). Single units were then split into pyramidal cell and interneurons based on their waveform duration, > 0.5ms and < 0.5ms respectively. A visualization of this classification is shown in Fig. S2A-C.

#### Measures of Intrinsic Firing Characteristics

Measures of intrinsic cell firing included average firing rate, bursting propensity (105), and coefficient of variation II (CV2) (40). Average firing rate was calculated by dividing the total number of spikes over the session duration. Bursting propensity was calculated from a neuron’s autocorrelogram (Fig. S2C) by dividing the peak value between 2*ms* – 9*ms* by the mean value between 40*ms* – 50*ms*. This value was then normalized by the peak when the peak value was greater than the baseline, or normalized by the baseline for peak values less than the baseline. This allowed values to range from −1 to 1, with higher positive values indicating higher bursting. CV2 was calculated as:

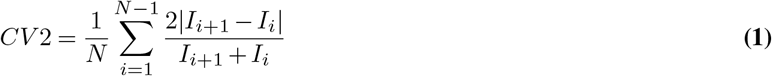

where *t* is the spike time stamp at index *i* and *N* is the total number of spike times. A CV2 of 1 indicates that a spike train is similar to a sequence of intervals generated by a Poisson process (40).

#### Sharp-wave ripple detection

Sharp-wave-ripples were detected similar to the methods used in previous studies (106). First, the local field potential (LFP) was extracted by re-sampling the wide-band signal from 32kHz to 1250Hz. Next, the LFP was bandpass-filtered (FIR filter: passband: 80-250Hz) (107) and the instantaneous amplitude was then extracted using a Hilbert transform. The instantaneous amplitude was then smoothed with a Gaussian filter (s.d.= 4ms) and Z-scored. Candidate ripple events were then detected where the peak amplitude was > 3 s.d. for ≥ 15ms and event edges were defined when the amplitude extended back to the mean; overlapping events were merged. Candidate events that occurred when the animal was moving (> 4 cm/sec) were then discarded. To remove false-positive SPW-Rs, candidate events were discarded if they co-occurred with moments of high amplitude (> 0.85(r)) intracranially derived electromyography (EMG). Finally, candidate events that did not co-occur with moments of highly synchronous multi-unit activity were discarded.

#### Intracranially derived electromyography

Although simultaneous EMG recordings were not taken, EMG from the neck, jaw, and face muscles was estimated from intracranially recorded LFP by detecting the zero time-lag correlation coefficients between 300–499Hz filtered signals recorded at different sites (108,109). Important to note, EMG was not estimated for animals implanted with the 8-tetrode bundle drive design as tetrodes in this design are close together and share similar LFP signals as a result. Pearson’s correlation coefficients were calculated for each pair of channels across a 25ms sliding window. The coefficients were then averaged which created an estimated EMG signal sampled at 40Hz. Intracranially derived and direct EMG recordings have been previously found to be highly correlated (108). Similarly, intracranial EMG is highly correlated with motion/accelerometer measures and chewing moments (109).

#### Multi-unit synchronous event detection

Synchronous epochs in multi-unit spiking were detected using a method similar to previous work (50). Spike times from all simultaneously recorded units were concatenated and binned into 1-ms windows and subsequently smoothed with a Gaussian kernel (s.d.= 15 ms). Epochs of synchronous spiking were identified as periods exceeding 3 s.d. above the mean. The edges of the event were defined as when the multi-unit activity returned to the mean. Epochs that occurred while the animal was moving (velocity of > 4 cm/sec) were then discarded. Epochs shorter than 15ms were also discarded.

#### SPW-R modulation of single units

Putative pyramidal cells and interneurons with at least 100 spikes in a given session were classified and analyzed according to their SPW-R modulation similar to previously used methods (35). First, cross-correlograms were constructed for each unit relative to all SPW-Rs detected in a session. For this, spikes were binned in 5ms bins with a ±500*ms* window around each SPW-R. To assess if a unit was significantly modulated by SPW-Rs, spike times were shuffled 400 times to create a null distribution where 95% upper and lower confidence intervals were calculated. Units that crossed either above or below the confidence intervals for at least a consecutive 20*ms* were classified as positively or negatively modulated. The participation probability of each unit was calculated by the number of times a unit fired at least one spike during a SPW-R divided by the total number of SPW-Rs detected in a session. As well, the fraction of units participating in each SPW-R was calculated as the number of units that fired at least one spike divided by the number of recorded units. Participation was only assessed in individual units that had ≥ 100 spikes and ≥ 50 SPW-Rs in a recording session.

#### Excitation-inhibition balance

To estimate the relative excitation-inhibition dynamics during SPW-R epochs, we monitored pyramidal cell and interneuron population activity over time. To do this, we binned spikes from pyramidal cells and interneurons separately into 10ms bins and smoothed each vector with a Gaussian kernel (s.d. 15ms). These two vectors were Z-scored in order to normalize the population firing around its baseline. This normalization controlled for instances where a recording session did not have the same number of pyramidal cells and interneurons and for inherit differences in firing characteristics between the two populations. Thus, any increases in activity during SPW-Rs were relative to the population’s baseline firing. This procedure is visualized in Fig. 4A. To assess the relative population activity, we identified the peak values from both vectors during each SPW-R (within-ripple pyramidal peak $ within-ripple interneuron peak). The within-ripple interneuron peak value was subtracted from the within-ripple pyramidal peak value for each SPW-R to estimate the relative difference between excitation-inhibition during SPW-Rs. Values above 0 indicated a bias towards excitation, whereas values below 0 indicated a bias towards inhibition.

#### Cell assembly analyses

Cell assemblies were identified as previously described (110). Significant co-firing patterns were detected using an unsupervised statistical method based on independent component analysis (ICA). Spike trains for each neuron were binned into 25ms time windows and z-score transformed to eliminate biases due to differences in average firing rates. Next, a principal component analysis was applied to the binned spike matrix (*Z*). The correlation matrix of *Z* was given by 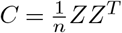 and eigenvalue decomposition of C was given by:

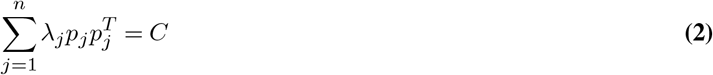

where *λ_j_* is the *j^th^* eigenvalue of *C* and *p_j_* is its corresponding eigenvector. The Marcenko-Pastur law was used to estimate the number of significant patterns embedded within *Z*. For a *nXB* matrix, an eigenvalue exceeding λ_*m*_*ax*, defined by 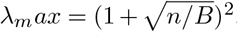, signifies that the pattern given by the corresponding principal component explains more correlation than would be expected if the neurons were independent of each other (111). The number of eigenvalues exceeding λ_*m*_*ax* was defined as *N_A_* and therefore represents the minimum number of distinct significant patterns in the data (110,112). The significant principal components were then projected back onto the binned spike data

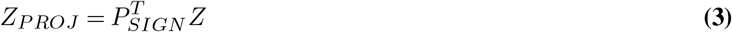

Where *P_SIGN_* is the *nXN_A_* matrix with the *N_A_* principal components as columns.

Independent component analysis (ICA), using the fast ICA algorithm (113), was then applied to the matrix *Z_PROJ_*. That is, an *N_A_XN_A_* unmixing matrix *W* was found such that the rows of the matrix *Y* = *W^T^Z_PROJ_* were as independent as possible. The unmixing matrix *W* was then used to derive each cell’s weight within each assembly *V* =*P_SIGN_W*.

To determine the strength of the expressed assemblies, we tracked each assembly pattern *v_k_* over time by:

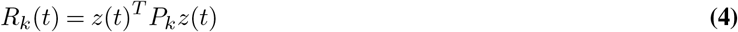

where *z*(*t*) is a smooth vector-function containing for each neuron its *z*-scored instantaneous firing-rate and *P_k_* is the matrix projecting *z*(*t*) to the activation-strength of the assembly pattern *k* at time *t*. To increase the temporal resolution, *z*(*t*) was obtained by convolving the spike-train of each neuron with a Gaussian kernel 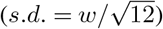 and then *z*-scoring each trace. *w* was set to 25ms to match the bin-size used to identify the assembly patterns. Assembly-activations were defined as peaks exceeding *R_THRES_* = 5.

Within ripple assembly strength was defined as the median peak activation value of events with R > 5 occurring during SPW-R epochs. Only sessions with ≥ 25 SPW-R events were considered for assembly analysis in order to assess distribution parameters.

#### Detection of awake replay

During pauses in exploration, ensembles of HPC neurons re-express firing sequences corresponding to the sequence of firing that occurred during recent navigation. These “replay” events co-occur with sharp-wave ripples and are associated with increased hippocampal-cortical communication (6–8). A Bayesian decoding approach (see (49) for original method) was used to detect and analyze replay events (see (50,51,54,55,114,115) for use cases of this method to investigate replay).

Spike counts from each multi-unit synchronous event were first binned into 20*ms* time bin *t* from *N* units *O_t_* = (*o*_1*t*_*o*_2*t*_…*o*_*Nt*_) (500*ms* time bins were used for active behavior decoding). Then the posterior probability distribution were calculated for each *t* over binned positions (3 cm) along the linear track using Bayes’ rule:

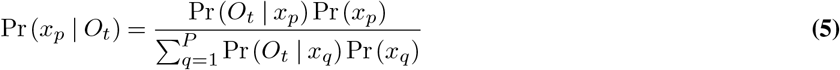

Where *x_p_* is the center of the *p* – *th* position bin. Because we assumed Poisson firing, the prior probability, *Pr*(*O_t_*|*x_p_*,), for the firing of each unit *n* is equal to

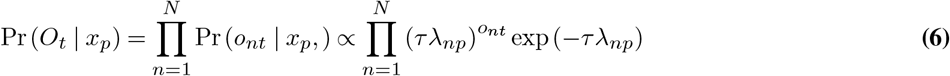

where *τ* is the duration of time (20 ms) and *λ_np_* is the mean firing rate of the *n* – *th* unit in the *p* – *th* position. We assumed a uniform prior distribution *Pr*(*x_p_*) over the position bins. This implementation of Bayesian decoding was carried out using tools in the Python package Nelpy (103).

Each replay candidate was considered a significant replay event if the series of decoded positions is more consistent with an ordered trajectory compared to a surrogate distribution. Linear regression was used to fit a line to the posterior probability distribution. A Bayesian replay score for a given event was defined as the sum probability mass under the fit line within a bandwidth (of 21 cm) (50). For each candidate event, we generated 1500 surrogates of the posterior probability distribution by circularly shifting each column of the posterior probability matrix by a random amount. This type of surrogate generation was specifically selected as it allows for the inclusion of stationary trajectory events, which have been described previously (116,117). A Monte Carlo p-value for each event was obtained from the number of surrogate events with replay scores higher than the observed score. The threshold for significance was held at 0.05. Replay events were identified from events with at least 10 active pyramidal cells with a peak firing rate of ≥ 1Hz and a peak to mean firing rate ratio of 1.5. As well, candidate events with many (> 50%) inactive spike count bins were discarded.

This Bayesian decoding algorithm was also used to estimate the animal’s location during active running (speed > 4 cm/s), similar to previous studies (50,118–120).

Recording sessions with poor decoding accuracy of the animal’s position were excluded from the replay analysis. Decoding accuracy was assessed with a surrogate analysis where the spike times from each unit were circularly shifted by a random amount. This was done in order to preserve the temporal components of each spike train but randomize the spike times relative to the position in which they fired. Linear regressions between the real and decoded position of the animals were calculated, and a Monte Carlo p-value for each recording session was obtained using the *R*^2^ values from each regression. Only sessions with a significant Monte Carlo p-value < 0.05 were used to quantify replay.

#### Statistical Analysis

Statistics were computed using custom software written in R. No data were excluded based on statistical tests. Using R packages lme4 (121) and lmerTest (122), linear mixed-effects models were used to assess group differences. The basic model used was *feature* ~ *group* + (1|*rat*/*session*). Normality and Independence of errors were inspected visually and corrected with an appropriate transform if the errors were skewed. Group differences in proportion were modeled using generalized linear models with mixed effects and fit using the glmer() function with family = “binomial”. The significance threshold was set at *α* = 0.05 unless otherwise stated.

---

**Figure S1.**
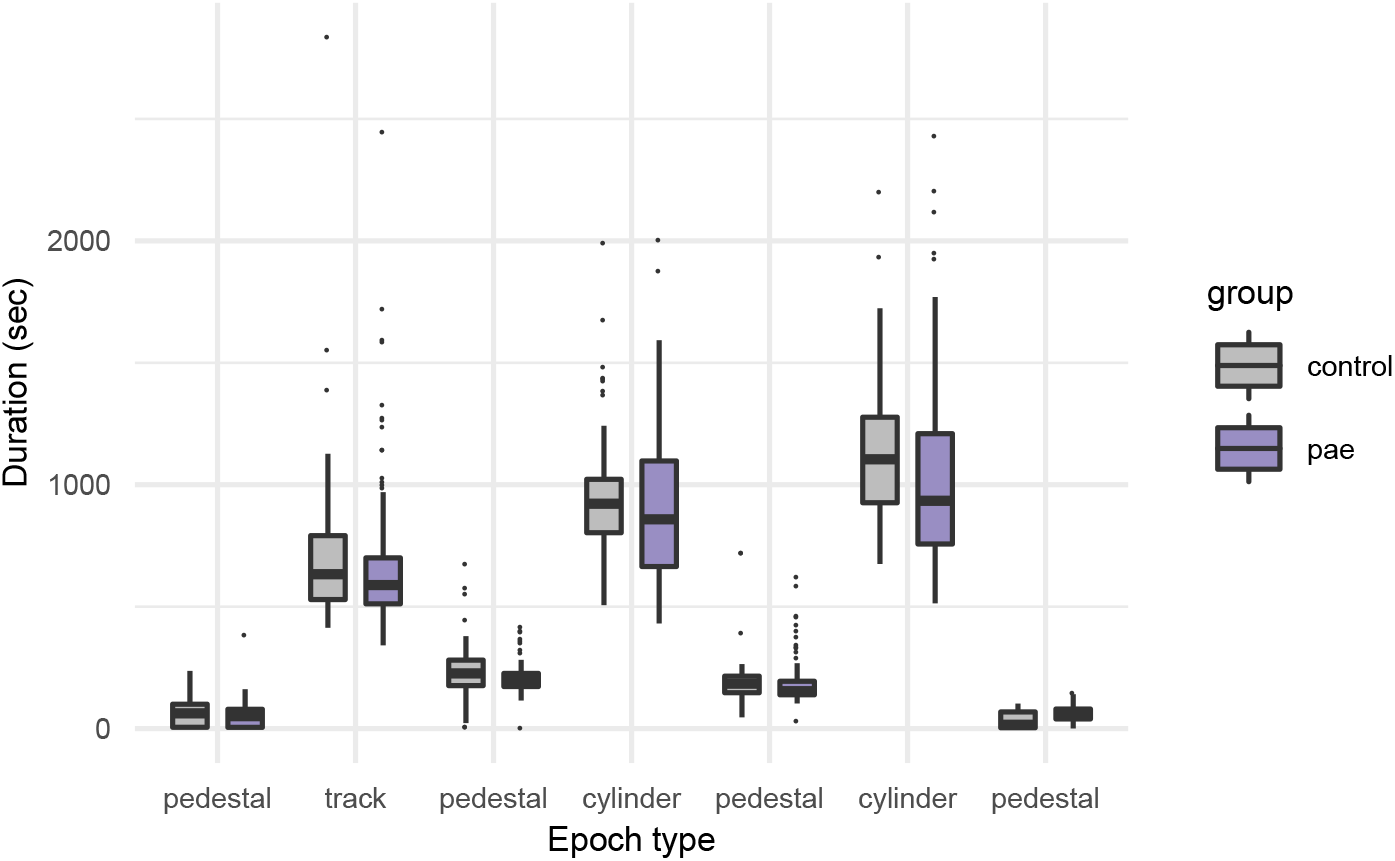
Epoch duration was not significantly different between groups across all epoch types (*ps* ≥ 0.3322).

**Figure S2.**
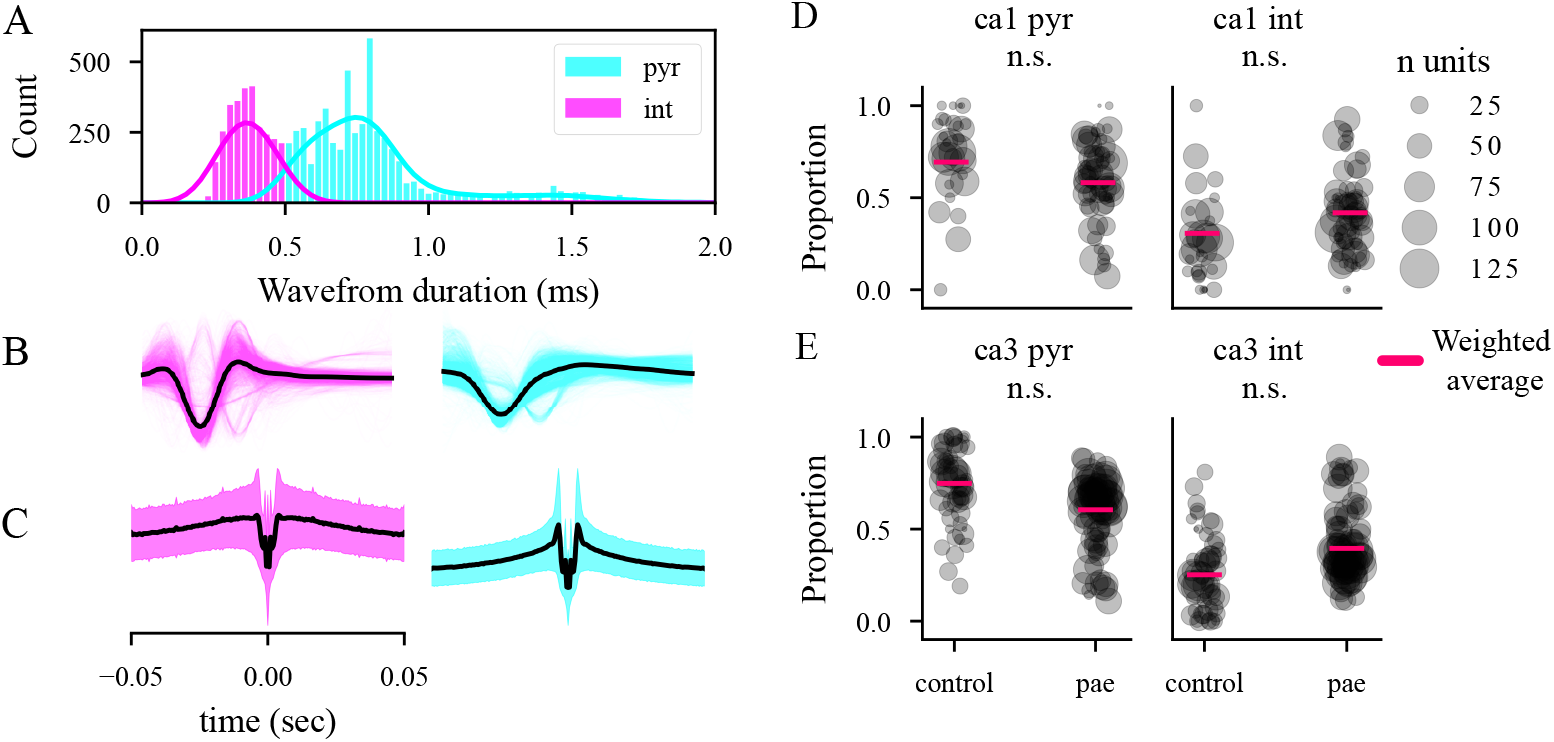
Physiological classification of pyramidal cell and interneurons. (A) Shows duration of the extracellular spike. Spike duration was estimated using the inverse peak frequency of a wavelet transform of the spike waveform. The distribution spike durations is split into two classes pyramidal cells (orange) with durations above 0.5ms and interneurons (blue) with durations below 0.5ms. (B) Shows the spike waveforms from all pyramidal cells and interneurons with the mean trace for each class shown in black. Individual waveforms have a transparency applied. (C) Shows the mean (black) and standard deviation (shading) spike autocorrelograms from all pyramidal cells and interneurons. Note that pyramidal cells discharge in bursts, which is visualized by the sharp peaks near the center of the autocorrelogram, whereas interneurons discharge in a more regular pattern. (D-E) Shows the proportion of cells per session classified as interneurons and pyramidal per group in HPC subregions CA1 & CA3. Group differences in proportions of classified units were compared with generalized linear models using rat and recording session as nesting factors. As such, data here is represented as a single point per recording session with the size of the point representative of the number of units recorded in that session. Weighted group averages are shown as pink lines over the scatted points. The weights in the weighted average are the number of units in each session.

**Figure S3.**
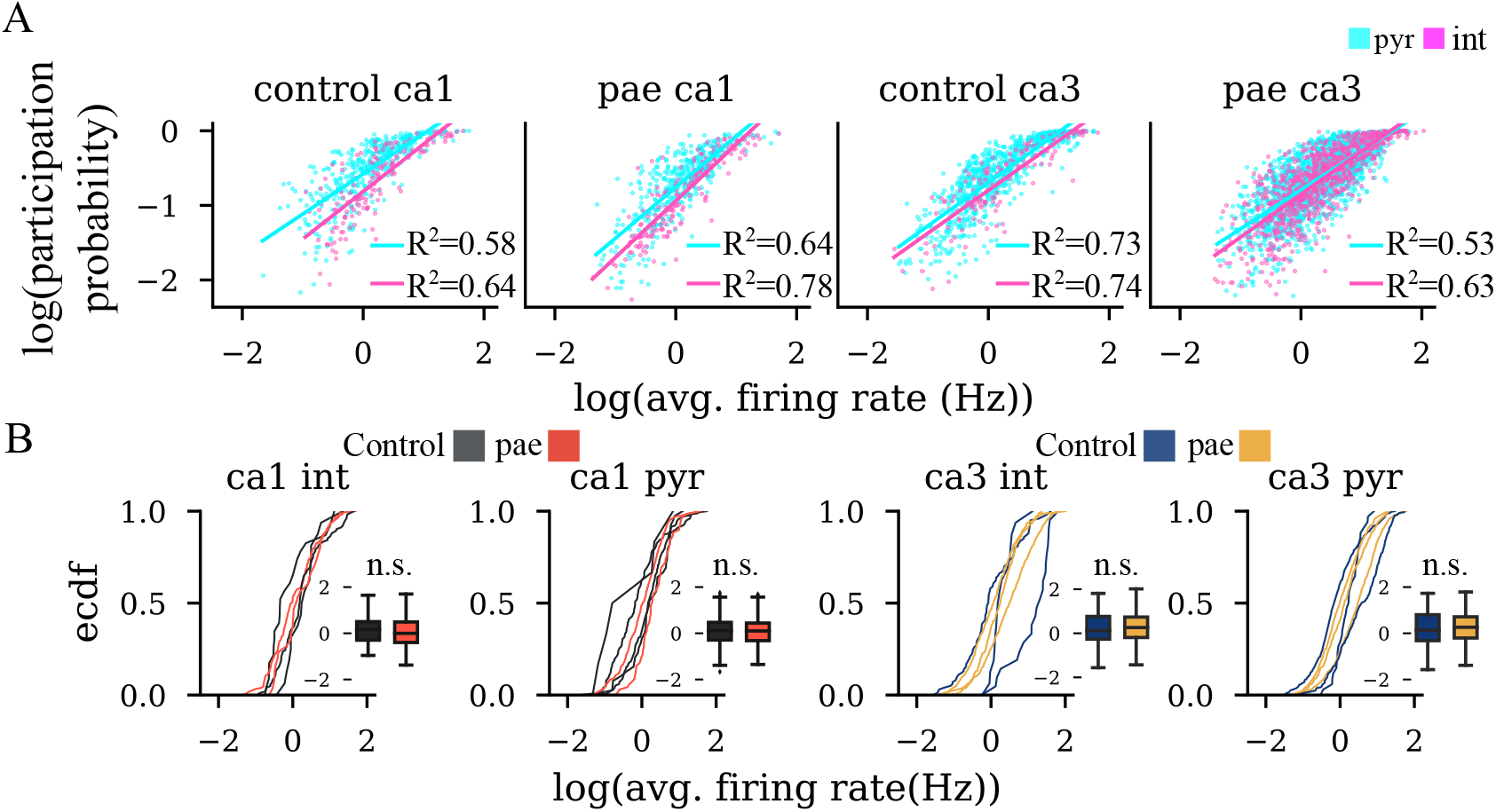
Average firing and SPW-R participation probability. (A) Shows the positive relationship between average firing rates and SPW-R participation probability between groups, subregions, and cell types. (B) A comparison of average firing rates between groups. No significant group difference was detected (*ps* ≥ 0.30).

**Figure S4.**
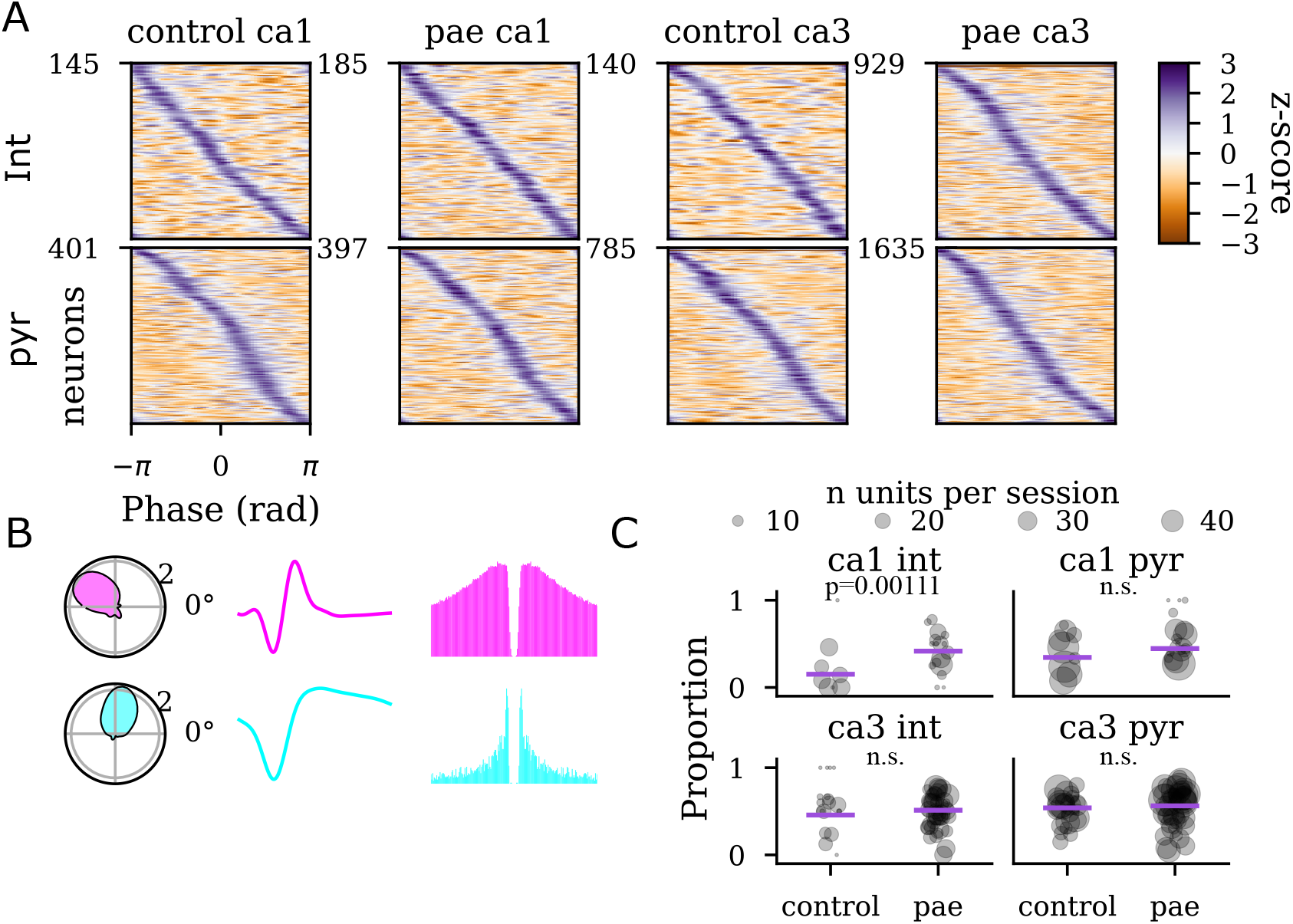
Phase dynamics of individual units during SPW-Rs. (A) Peri-SPW-R (±100*ms*) Z-scored phase plots demonstrating how individual units are modulated by SPW-R phase. Neurons are organized by the phase that maximized the firing rate of each unit. (B) Phase polar plots, average waveforms, and autocorrelograms of two control CA1 example cells (top: interneuron, bottom: pyramidal cell) showing strong phase modulation to SPW-Rs. (C) Proportion plots showing the proportion of SPW-R phase modulated units. Group differences in proportions of significant SPW-R phase modulated units, accessed with having a Rayleigh p-value below 0.05, were compared with generalized linear models using rat and recording session as nesting factors. As such, data here is represented as a single point per recording session with the size of the point representative of the number of units recorded in that session. Weighted group averages are shown as purple lines over the scatted points. The weights in the weighted average are the number of units in each session. Group weighted average proportions are listed: CA1 pyramidal: control 34%, PAE 45%; CA1 interneuron: control 15%, PAE 42%; CA3 pyramidal: control 54%, PAE 56%; CA3 interneuron: control 46%, PAE 51%

